# Retinol Binding Protein 4 reactivates latent HIV-1 via the JAK/STAT5 and JNK pathways

**DOI:** 10.1101/2025.02.26.640366

**Authors:** Chiara Pastorio, Khumoekae Richard, Manuel Hayn, Lennart Koepke, Andrea Preising, Nico Preising, Ludger Ständker, Matthew Fair, Jessicamarie Morris, Emmanouil Papasavvas, Honghong Sun, Armando Rodríguez, Karam Mounzer, Sebastian Wiese, Pablo Tebas, Yangzhu Du, Gregory M. Laird, Konstantin M.J. Sparrer, Luis J. Montaner, Frank Kirchhoff

## Abstract

Reactivation of the latent viral reservoirs is crucial for a cure of HIV/AIDS. However, current latency reversing agents are inefficient and the endogenous factors that have the potential to reactivate HIV *in vivo* remain poorly understood. To identify natural activators of latent HIV-1, we screened a comprehensive peptide/protein library derived from human hemofiltrate, representing the entire blood peptidome, using J-Lat cell lines harboring transcriptionally silent HIV-1 GFP reporter viruses. Fractions potently reactivating HIV-1 from latency contained human Retinol Binding Protein 4 (RBP4), the carrier of retinol (vitamin A). We found that retinol-bound holo-RBP4 but not retinol-free apo-RBP4 strongly reactivates HIV-1 in a variety of latently infected T cell lines. Functional analysis revealed that this reactivation depends on the JAK/STAT5 and JNK pathways but does not require retinoic acid production. High levels of RBP4 were detected in plasma from both healthy individuals and people living with HIV-1. Physiological concentrations of RBP4 induced significant viral reactivation in latently infected cells from individuals on long-term antiretroviral therapy with undetectable viral loads. As a potent natural HIV-1 latency-reversing agent, RBP4 offers a novel approach to activating the latent reservoirs and bringing us closer to a cure.

## INTRODUCTION

HIV-1 is the main causative agent of AIDS, a pandemic that has already affected more than 80 million people and led to an estimated 42.3 million deaths as of 2024 (Global HIV Statistics 2024, UNAIDS). HIV-1 infections are characterized by progressive depletion of CD4+ T-helper cells and chronic immune activation.^1^ Combined antiretroviral therapy (cART) prevents HIV-1 replication and has greatly improved the life expectancy and quality of infected individuals.^2^ However, cART fails to cure HIV infection and treatment is required for life. A reason for this is the livelong persistence of the virus in a small percentage of resting CD4 + T cells even under effective cART.^3,4^ These cells carry stably integrated and transcriptionally silent but still replication-competent proviruses in a reversibly non-productive state of infection. While they do not produce virus particles in this resting state, they can be reactivated to produce virions that may cause spreading infection and disease upon treatment interruption.^5,6^ Therefore, targeting and eradicating the latent viral reservoirs is essential for achieving a cure for HIV/AIDS.

One therapeutic strategy to reduce HIV-1 reservoir size is the so-called “kick and kill” approach.^7,8^ The first step is the administration of latency reversing agents (LRAs) to patients on cART. Reactivated cells start producing virions and become susceptible to antiretroviral drugs and host cytolytic effector mechanisms.^9^ Several compounds have been characterized for their ability to induce reactivation of HIV-1 gene expression from latency including protein kinase C (PKC) agonists, histone deacetylase inhibitors (HDACi), histone methylation inhibitors (HMTi), or DNA methyltransferase inhibitors.^9^ Current approaches, however, have proven too ineffective to clearly reduce the number of latently HIV-infected cells.^10^

Upon interruption of cART, HIV-1 rebounds to detectable levels in most people living with HIV (PLWH) within a few weeks.^11^ It is currently poorly understood which endogenous factors reactivate the virus from latency. Their identification may not only help to better understand the mechanisms underlying viral rebound but also provide novel means to drive HIV-1 out of latency and reduce the size of the latent viral reservoirs. To address this, we performed a systematic screen of a hemofiltrate-derived peptide library for endogenous circulating activators of latent HIVs. We discovered that human Retinol Binding Protein 4 (RBP4) carrying retinol (vitamin A) strongly activates latent HIV-1 in various J-Lat cell lines. In agreement with a relevant role *in vivo*, RBP4 also reactivates HIV-1 in cells obtained from people living with HIV (PLWH) after long-term antiretroviral therapy. Our data support that the abundant carrier of retinol RBP4 promotes HIV-1 expression and offers prospects for reactivation of the latent viral reservoirs.

## RESULTS

### Identification of RBP4 as activator of latent HIV-1 proviruses

To identify endogenous circulating factors that reactivate latent HIV-1, we generated a protein/peptide library from human hemofiltrate (HF) using cation exchange (generating 8 pH pools) and reversed phase (RP) chromatography, resulting in 48 fractions per pool.^12^ As previously reported^12^, these libraries contain essentially all peptides and small proteins circulating in human blood in a lyophilized, bioactive and highly concentrated form. The total of 384 peptide-containing RP-HPLC fractions from the HF peptide library were analysed for factors reactivating latent proviruses (i.e. are inducing GFP expression) in J-Lat 11.1 cells, which carry an *env*-defective HIV-1 construct containing the *GFP* reporter gene in place of the accessory *nef* gene.^13^ Phorbol myristate acetate (PMA) and tumor necrosis factor-alpha (TNF-α), two well-established activators of HIV-1 latency, were used as positive controls. A strikingly high number of HF fractions from different pH pools efficiently reactivated HIV-1 transcription (Fig. 1a). In several cases, the levels of GFP expression were about as high as those achieved with PMA and TNF-α (example shown in Fig. 1b), which are highly potent but impractical for therapeutic use due to high toxicity and nonspecific activation of immune cells. The most active fractions were combined and further separated by additional chromatographic steps (Supplementary Fig. 1a). SDS-PAGE revealed that the active fractions contained a protein of about 21 kDa (Supplementary Fig. 1b). MALDI-TOF MS analysis of the activating fraction F42 obtained after the final purification step showed the mono, double and triple charged species of Retinol Binding Protein 4 (RBP4) (Fig. 1a; Supplementary Fig. 1c, d, e), a member of the lipocalin family^14^. Specifically, the species identified (m/z 20960.76) corresponds to RBP4 lacking the C-terminal leucine (RBP4 1-182). Notably, forms of RBP4 lacking the C-terminal leucine have been previously detected in serum of hemolysis patients.^15^ The N-terminal secretory signal peptide of 18 amino acids (aa) is cleaved upon processing and RBP4 circulates as a protein of 183 amino acids in human blood.^16^ RBP4 is the main carrier of Vitamin A (Retinol) and transports it to different target tissues.^17^ In addition, RBP4 contributes to insulin resistance in obesity and type 2 diabetes and may induce TLR2/4 signalling in immune cells.^17,18^ RBP4 was readily detectable in active HF fractions (Fig. 1c) and its concentrations correlated with the levels of HIV-1 reactivation indicated by the percentages of GFP+ cells (R^2^ = 0.667; P<0.0001). Thus, our results identified RBP4 as potent circulating activator of latent HIV-1 proviruses.

**Fig. 1.**
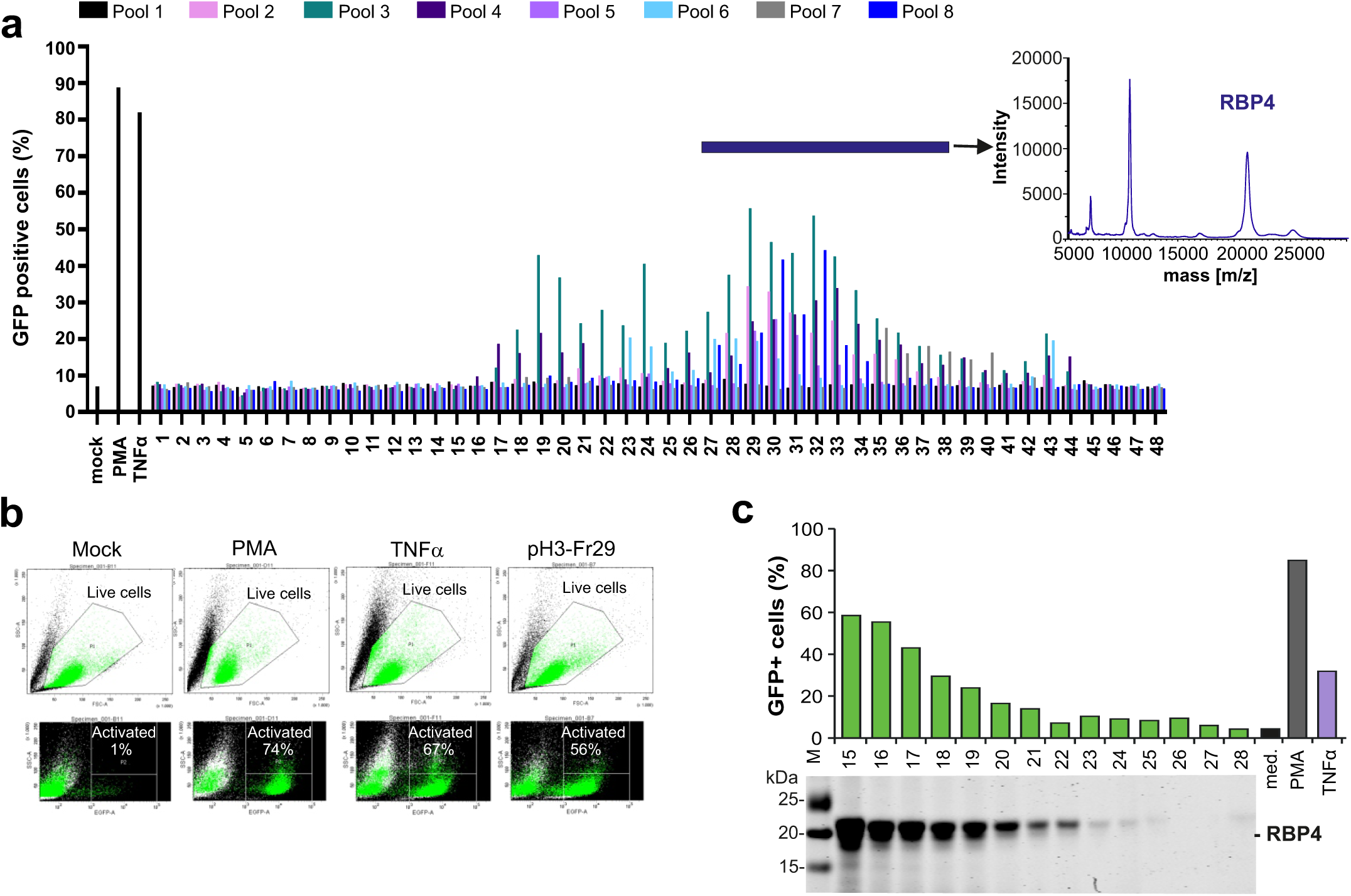
Identification of RBP4 as enhancer of latent HIV. (**a**) (Left panel) Activation of latent HIV-1 eGFP reporter proviruses in J-Lat 11.1 cells treated with peptide fractions of a hemofiltrate (HF)-derived library. The color of the bar indicates to which pool of the peptide library the fraction belongs. Mock indicates the absence of peptide fractions; PMA and TNFα were used as positive controls. Values represent the mean ± SD of triplicate infections. Fractions highlighted by the upper bar were used for further purification. (Right panel) MALDI-TOF spectrum of RBP4 shows a dominant signal of m/z 20960.75, which closely matches the theoretical m/z value of the RPB4 (1-182, 20959.42 Da) truncated variant lacking the C-terminal leucin. (**b**) Example for activation of latent HIV-1 in J-Lat 11.1 cells by a single HF-derived fraction compared to the PMA and TNFα controls, determined by flow cytometry. (**c**) The levels of RBP4 in HF fractions detected by western blot after purification (lower panel) correlate with the percentages of cells showing reactivation of HIV-1 eGFP reporter proviruses.

### RBP4 activates latent HIV-1 in a variety of T cell lines

Our initial screen was performed using the J-Lat 11.1 subclone that is highly responsive to agents inducing latent HIV-1 reactivation.^13,19^ A variety of J-Lat cell lines harbouring either proviral sequences containing GFP in place of *nef* or LTR-Tat-IRES-GFP expression cassettes thought to be silenced by different mechanisms are available.^13^ To assess the spectrum of latency reversal activity, we examined the effect of RBP4 on a total of 10 different J-Lat subclones. RBP4 dose-dependently and efficiently reactivated latent HIV-1 proviruses in the 10.6 and 11.1 cell lines but had no or only modest effects in the J-Lat 6.3, 8.4, 9.2 and 15.4 lines (Fig. 2a). Similarly, RBP4 enhanced GFP expression in the A1, A2 and H2 J-Lat Tat GFP reporter cell lines, while little induction of GFP expression was observed in A7 J-Lat cells (Fig. 2b). In addition, RBP4 also efficiently enhanced p24 production in the Jurkat J1.1 T cell clone that harbours the HIV-1 LAV-1 strain and has defects in CD3 signalling (Fig. 2c).^20^ In contrast, RBP4 did not significantly stimulate p24 antigen expression in the chronically HIV-1 LAV-1-infected promonocytic U937 cell line derivative U1 and in the HIV-1 LAV-1 latently infected lymphocytic CEM cell line ACH-2 (Fig. 2c).^21^ In most cases, latently infected cells that strongly responded to PMA also showed efficient latency reactivation in response to treatment with TNFα and RBP4. However, in J-Lat A1 and U937 U1 cells PMA was substantially more efficient than TNFα or RBP4 (Fig. 2). Overall, RBP4 induced HIV-1 reactivation from latency to similar levels as PMA or TNFα in several of the model cell lines tested. Differences in proviral integration sites, chromatin context, and transcription factor dynamics are known to significantly affect the responsiveness of specific J-Lat cell lines to stimulation by PMA and TNFα, and this was also observed with RBP4.

**Fig. 2.**
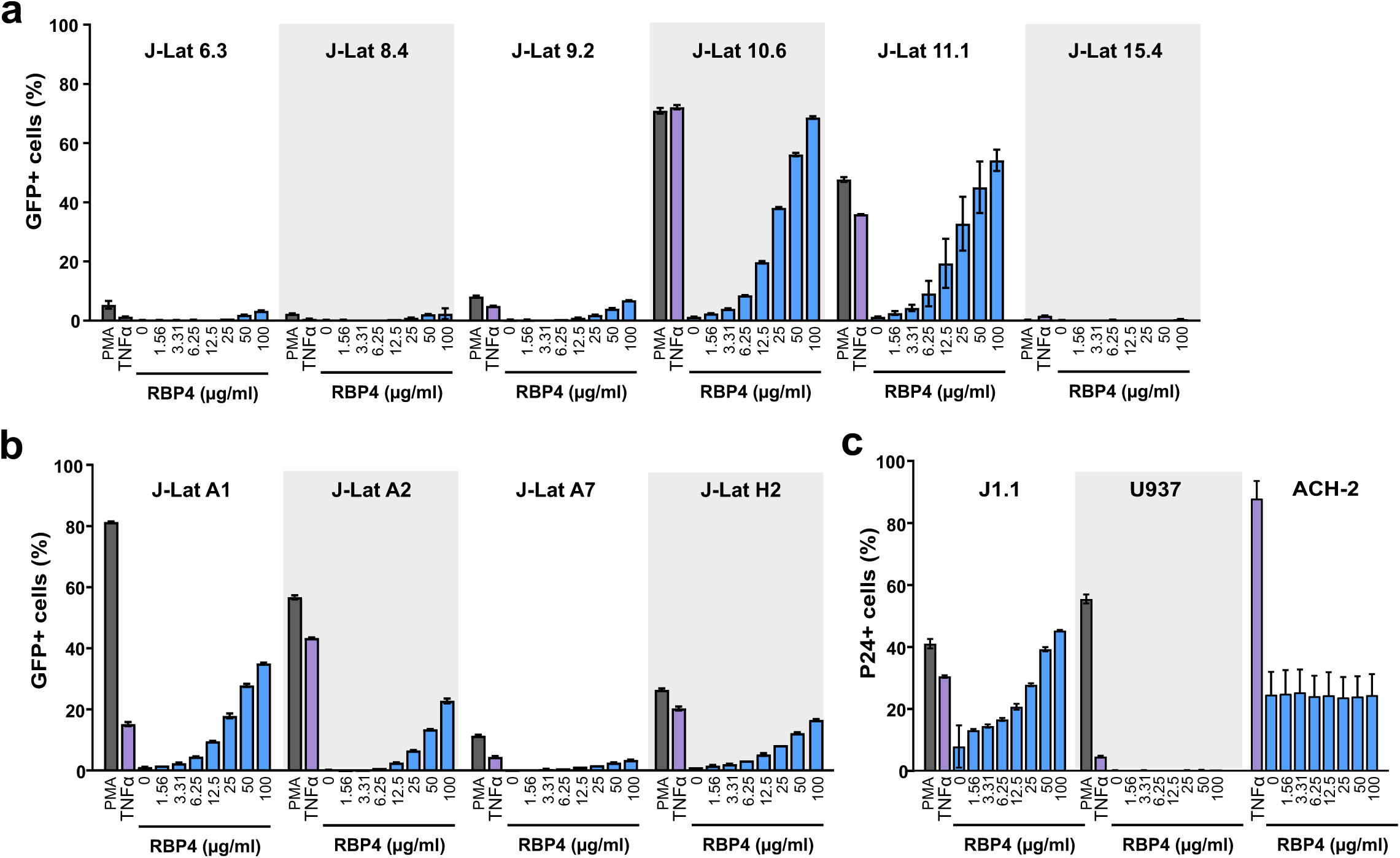
RBP4-mediated reactivation of latent HIV-1 in various cell lines. (**a, b**) J-Lat cell lines harboring (a) full-length HIV-1 eGFP proviruses or (b) LTR-Tat-IRES-GFP cassettes were treated with the indicated concentrations of RBP4 or PMA and TNFα for control and the percentages of green cells were determined by flow cytometry. (**c**) Jurkat J1.1 cells, U937 and ACH-2 cells harboring full-length intact HIV-1 proviruses were treated as in panels A and B but reactivation was determined by intracellular p24 antigen staining.

### Retinol is required but not sufficient for HIV-1 reactivation

To obtain further insights into its latency reversing effects, we analysed RBP4 purified from four different sources, i.e. hemofiltrate, blood plasma, urine or recombinant expression. We found that RBP4 purified from hemofiltrate or blood plasma efficiently reactivated latent HIV-1 in J-Lat 10.6 cells (Fig. 3a). In contrast, recombinant RBP4, as well as RBP4 isolated from human urine, displayed little if any activity although proteins of correct size were readily detectable by western blot analysis (Fig. 3b). In agreement with published data^22^, the levels of RBP4 in plasma from eight healthy individuals determined by ELISA varied between 20 and 40 µg/ml (Fig. 3c). These concentrations were well sufficient to reactivate latent HIV-1 in a variety of model cell lines (Fig. 2). Thus, RBP4 is capable of reactivating HIV-1 from latency at physiological concentrations.

**Fig. 3.**
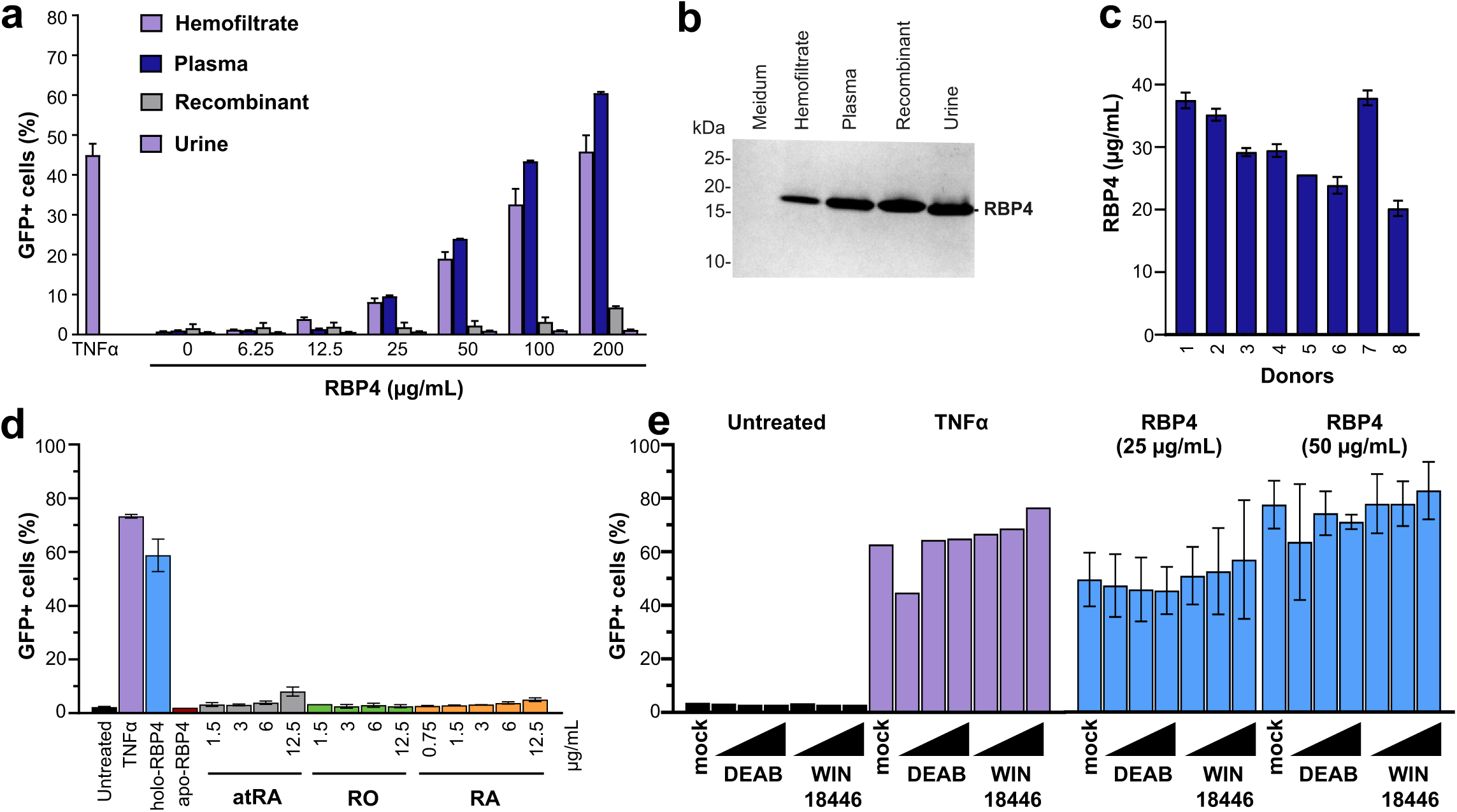
Impact of RBP4 from different sources and free retinol and its metabolic products on HIV-1 reactivation. (**a**) RBP4 from the indicated sources was tested for its ability to reactivate latent HIV-1 in the J-Lat 10.6 cell line at the indicated concentrations and examined by western blot analysis and (**b**). TNFα was used as positive control. (**c**) Concentration of RBP4 in the plasma from eight healthy donors measured by ELISA assay. (**d**) J-Lat 10.6 cells were treated with the indicated concentrations of all trans-retinal (atRA), retinol (RO) and retinoic acid (RA) and the percentage of GFP positive cells was determined by flow cytometry. TNFα, holo-RBP4 and apo-RBP4 are shown for comparison. (**e**) J-Lat 10.6 cells were pretreated with DEAB or WIN18,446 inhibiting conversion of retinal to retinoic acid and subsequently left untreated or treated with TNFα or the indicated concentrations of RBP4. Activation of latent HIV-1 proviruses (eGFP) was measured by flow cytometry.

The main function of RBP4 is to transport retinol (vitamin A) from the liver to peripheral tissues. Two different isoforms of RBP4 have been reported to exist in serum, namely holo-RBP4 (RBP4 bound to retinol) and apo-RBP4 (Retinol-free RBP4), that remains in the blood after the release of its cargo into the target cells.^22^ Since retinol has a very low molecular weight (0.286 kDa), these two forms migrate at similar size in western blots (Fig. 3b). It is known, however, that ∼90% of RBP4 circulating in the blood stream contains retinol.^23^ RBP4 largely circulates in complex with transthyretin (TTR) to prevent its loss from the circulation by filtration in the renal glomeruli.^24^ After delivery of vitamin A to the target tissue, RBP4 loses the affinity for TTR and apo-RBP4 is filtered through the glomeruli of the kidneys and ends up in urine. Thus, while RBP4 mainly exists in retinol-associated form in blood, it is retinol-free in urine.^24,25^ Similarly, recombinantly produced RBP4 lacks retinol. Thus, our results strongly suggest that retinol-free apo-RBP4 lacks the ability to reactivate HIV from latency.

It has been reported that retinoic acid (RA), the biologically active metabolite of retinol, reactivates latent HIV-1 proviruses.^26,27^ To determine whether retinol or its metabolites may not only be required but also sufficient for reactivation, we examined the effects of purified free forms of retinoids. All trans-retinal, retinol and retinoic acid alone all had little if any effect on HIV-1 latency reactivation in J-Lat 10.6 cells, while holo-RBP4 was highly effective (Fig. 3d). In addition, all trans-retinol was cytotoxic at higher concentrations (Supplementary Fig. 2a). Uptake of free unbound RA is typically minimal and under physiological conditions most RA in cells is generated from the delivery of retinol by RBP4 upon binding of the receptor ’stimulated by retinoic acid 6’ (STRA6) and its subsequent conversion to RA within the cells. To exclude that poor uptake was the reason for lack of proviral reactivation by free retinol and RA, we treated J-Lat 10.6 cells with DEAB or WIN18,446, which inhibit the conversion of retinal to retinoic acid upon delivery by holo-RBP4.^28^ Both agents had no inhibitory effect on reactivation of HIV-1 by TNF-α or RBP4 (Fig. 3e, supplementary Fig. S2b). Altogether, these results provide strong evidence that HIV-1 reactivation requires holo-RBP4, but is independent of retinoic acid production in the target cells.

### RBP4-induced HIV-1 reactivation involves both JAK/STAT5 and JNK signaling

Holo-RBP4 not only delivers retinol to cells but also activates JAK/STAT, as well as TLR4/JNK signaling.^29^ Both of these pathways play roles in HIV-1 transcription and it has been reported that the JAK1/2 inhibitors suppress HIV-1 reactivation from latent cells.^30^ Treatment with a STAT5 inhibitor (Biomol) significantly reduced RBP4-mediated reactivation of HIV-1 from latency in J-Lat 10.6 and 11.1 cells in a dose-dependent manner without causing toxic effects (Fig. 4a, supplementary Fig. 3a). JAK/STAT activation is mediated by the binding of holo-RBP4 to the STRA6 receptor and thought to involve transport of retinol.^31^ However, RBP4 has also been shown to activate JNK signaling via interaction with TLR4 without transporting retinol into the cells.^32^ JNK signaling promotes T cell activation and it has been reported that inhibition of the JNK/AP1 pathway promotes HIV-1 latency.^33^ In agreement with this, treatment with a JNK inhibitor dose-dependently decreased RBP4-mediated reactivation of latent HIV-1 in J-Lat 10.6 and J-Lat 11.1 cells by up to 70% (Fig. 4b, supplementary Fig. 3b). Since holo-RBP4 has been reported to exert complex effects on cell metabolism and signaling^34^, we also examined inhibitors of PKC (Rottlerin), ERK1/2 (U0126) and MEK1/2 (Selumetinib) signaling. However, none of these agents significantly affected HIV-1 reactivation in J-Lat 10.6 cells (Supplementary Fig. 4). In contrast, combined treatment with a STAT5 and JNK inhibitor strongly reduced the induction of proviral gene expression by RBP4, but had only modest effects on TNFα-induced reactivation (Fig. 4c). Altogether, these results show that both JAK/STAT5 as well as TLR4/JNK signaling contribute the RBP4-induced reactivation of latent HIV-1 in J-Lat cells.

**Fig. 4.**
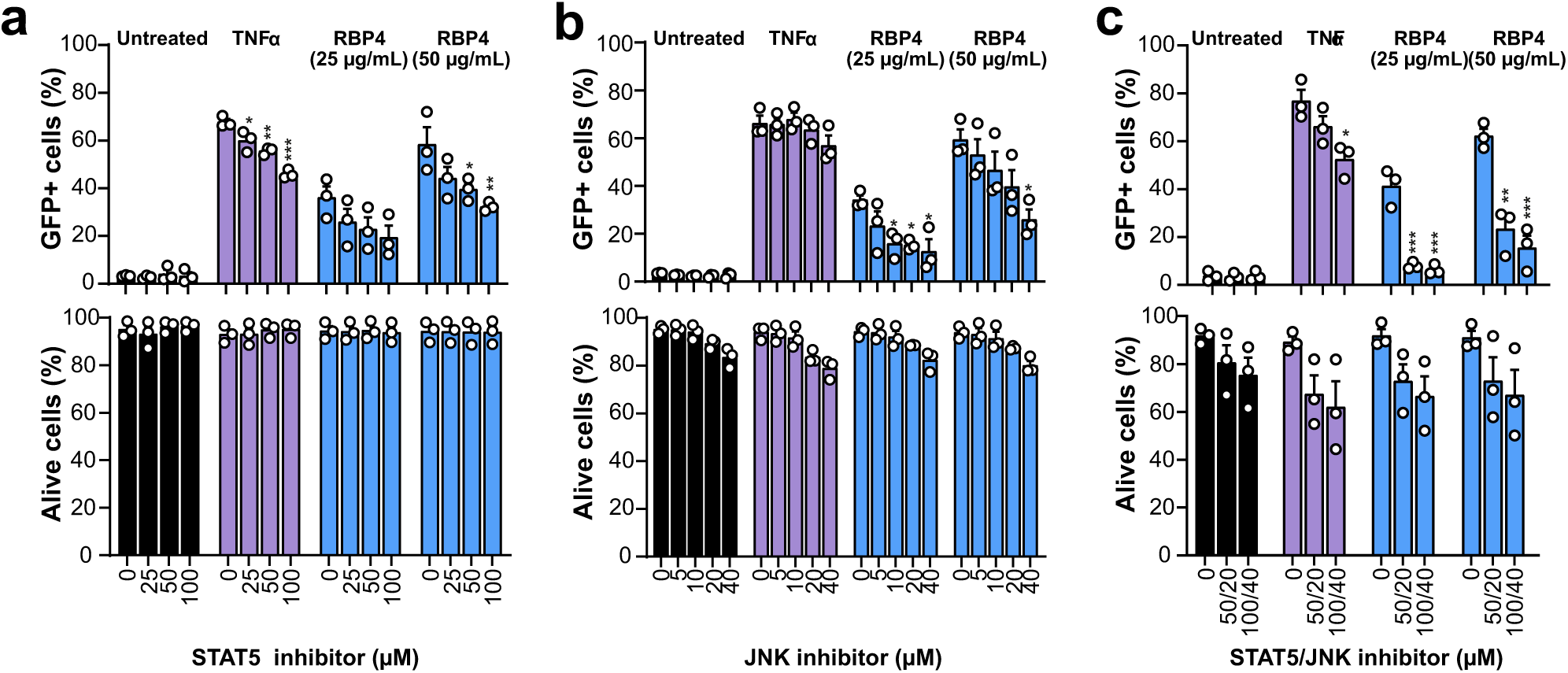
Role of JAK/STAT and TLR/JNK activation in RBP4-mediated HIV reactivation. J-Lat 10.6 cells were pretreated with the indicated concentrations of STAT5 (**a**) or JNK (**b**) inhibitor or their combination (**c**). Four hours later, the cells were left untreated or treated with TNFα or the indicated concentrations of RBP4. The percentage of cells showing reactivation of HIV-1 eGFP reporter proviruses (upper panel) and the cell viability (lower panel) were determined by flow cytometry.

To identify potential reasons for the differential responsiveness of the different J-Lat cell lines to RBP4, we determined the expression levels of its primary receptor STRA6.^35^ J-Lat cells that were readily reactivated by RBP4 (10.6 and 11.1) and clones that did not respond (8.4 and 9.2), expressed similar levels of STRA6 mRNA. In addition, primary activated CD4+ T cells just express low or no levels of STRA6 mRNA (Supplementary Fig. 5a). Moreover, RBP4 treatment did not induce expression of IL-6, IL-1ß or STRA6 receptor in the J-Lat cell lines irrespectively of HIV-1 reactivation (Supplementary Fig. 5b, 5c). It has been previously shown that differences in the responsiveness of different J-Lat cells lines are primarily linked to proviral integration sites and chromatin context. Altogether, our results imply that these features rather than differences in STRA6 expression levels also govern HIV-1 reactivation in J-Lat sublines upon RBP4 treatment.

### RBP4 reactivates latently infected cells derived from PLWH under ART

The levels of RBP4 are increased in various inflammatory diseases and inflammatory mediators are frequently enhanced in PLWH even after long-term ART. To determine the levels of RBP4 and assess potential correlations with the size of the latent viral reservoirs, we analyzed these parameters in a total of 64 ART-treated PLWH. The RBP4 levels ranged from about 10 to 50 µg per ml in plasma (Fig. 5a), which agrees with the circulating concentrations reported for healthy adults^36^. They did not correlate with the numbers of total, defective or intact proviruses in the corresponding individuals (Fig. 5a). However, the average concentrations ((26.25±8.68 µg/ml) were well sufficient for significant reactivation of HIV-1 in *in vitro* latency models.

**Fig. 5.**
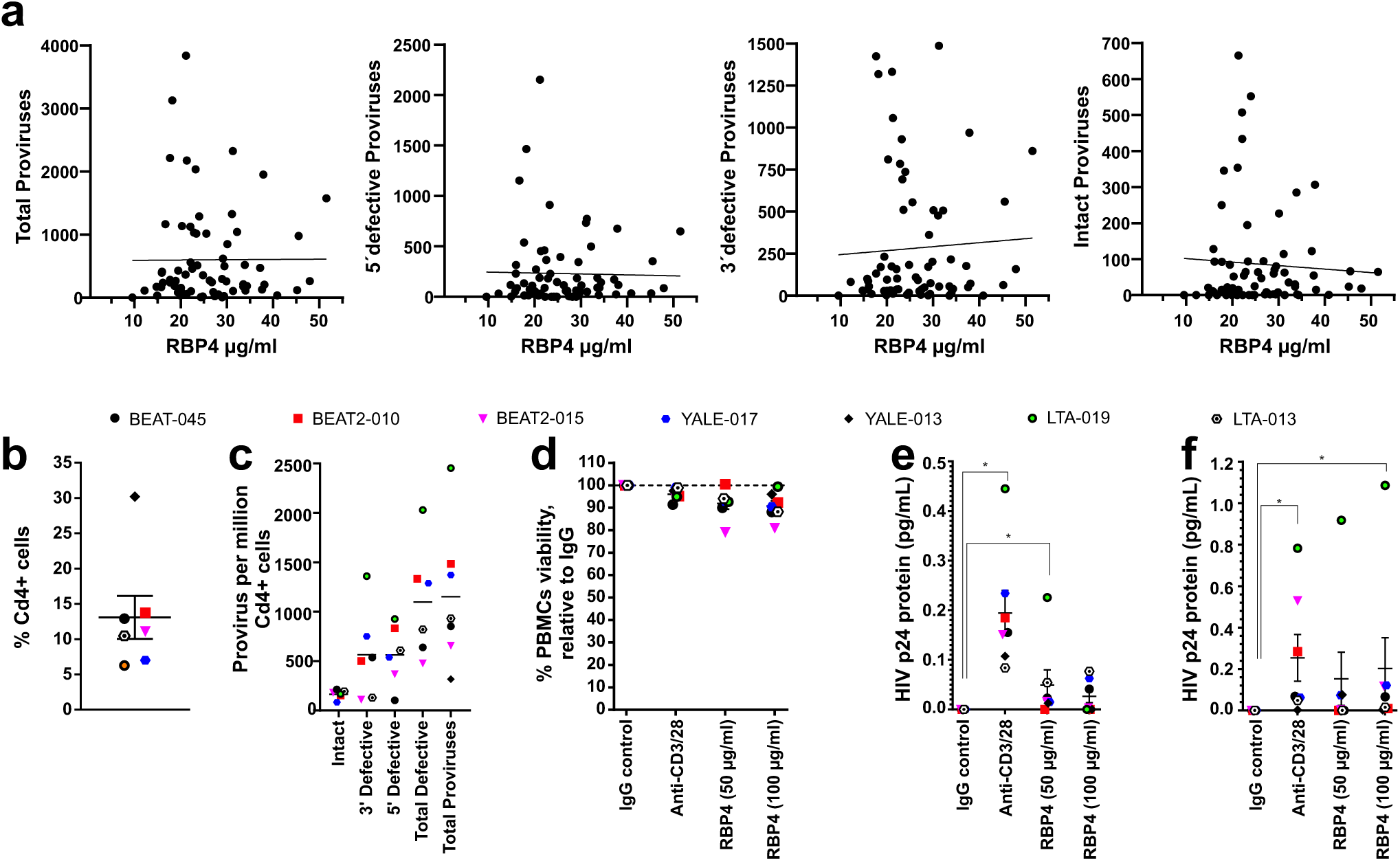
Reactivation of latent HIV-1 in PBMCs from PLWH under cART. (**a**) Correlation between the numbers of total, defective and intact proviruses and RBP4 plasma levels in the corresponding PLWH using Spearman correlation test. (**b**) Percentage of CD4+ cells of PBMCs from 7 PLWH under cART. (**c**) Percentage of intact and defective proviruses in CD4+ cells from cART-suppressed study participants, measured through IPDA. (**d**) Percent PBMC viability relative to IgG control after 72 hours treatment with LRAs. (**e, f**) Detection of Gag-p24 HIV-1 protein in cART suppressed PBMC cell pellets (e) and culture supernatants (f) after 72 hours treatment with LRAs as measured by SIMOA. *, p < 0.05 is shown as measured by Wilcoxon Signed-Rank test.

To assess the potential relevance of RBP4 expression *in vivo*, we examined its effect on PBMCs from seven people living with HIV (PLWH). All study participants were male and all but one African Americans (Table 1). The majority of them were under effective ART for more than 10 years and had undetectable viral loads at the day of blood collection. CD4+ T cells were in the normal range (Table 1, Fig. 5b). Using the well-established Intact Proviral DNA Assay^37^, we also assessed the composition and levels of intact and defective proviruses in CD4+ T cells from each donor. Proviral sequences were detectable in all individuals, but their levels varied substantially (Fig. 5c), with the frequency of intact HIV-1 proviruses being generally lower than the defective proviruses consistent with previous data^38^. To measure *ex vivo* latency reversal activities of RBP4, we next treated the PBMCs from the seven virally suppressed study participants with RBP4 (50 and 100 µg/mL), alongside anti-CD3/CD28 beads (50 µg/mL) as a positive control and Immunoglobulin G (IgG) (50 µg/mL) as the negative control for 72 hours, and measured the expression of HIV Gag-p24 protein in the cell pellets and culture supernatants using the ultra-sensitive Single Molecule Array (SIMOA)^39^. RBP4 was generally tolerated by the cells as no extensive toxicity was detected relative to IgG vehicle control (Fig. 5d). Our results show that RBP4 significantly reactivates latent HIV-1 in PBMCs from cART-suppressed individuals (p<0.05), even though its potency was inferior to the anti-CD3/28 positive control (Figs. 5e, 5f). IPDA analyses indicate that induction of p24 expression by RBP4 correlated with the number of total and defective but not intact proviruses (Supplementary Fig. 6). Altogether, the results show that RBP4 has a significant impact on HIV latency reactivation in cells from PLWH under effective ART at physiological concentrations.

**Table 1:** Study Participants baseline characteristics.

| Patient ID | Gender | Age | Ethnicity | Race | Date of HIV Diagnosis (mm/yyyy) | Date of starting ART (mm/yyyy) | Years of documented continual viral suppression | CD4 nadir recorded at first diagnosis | Curent ART regimen | CD4 Count on day of collection | Viral load Count on day of collection |
| --- | --- | --- | --- | --- | --- | --- | --- | --- | --- | --- | --- |
| BEAT-045 | Male | 45 | Not Hispanic | AA | 12/1998 | 2004 | 11 | 77 | Triumeq | 573 | <20 |
| BEAT2-010 | Male | 60 | Not Hispanic | White | 1999 | 2000 | 2 | Unk, >200 | Biktarvy | 1219 | <20 |
| BEAT2-015 | Male | 62 | Not Hispanic | AA | unk/1996 | 06/2003 | 12 | Unk, >200 | Triumeq | 396 | <20 |
| Yale-017 | Male | 43 | Not Hispanic | AA | 05/2003 | 2003 to 2006 | 12 | 349 | Symtuza | 653 | 51 |
| YALE-013 | Male | 57 | Not Hispanic | AA | 01/2002 | 2004 | 17 | 425 | Biktarvy | 694 | <20 |
| LTA-019 | Male | 72 | Not Hispanic | AA | 1988 | 10/2003 | 18 | 126 | Biktarvy | 921 | <20 |
| LTA-013 | Male | 56 | Not Hispanic | AA | 1989 | 10/2010 | 13 | 372 | Biktarvy | 411 | <20 |

## DISCUSSION

The persistence of HIV-1 in transcriptionally silent latent form in long-living memory CD4+ T cells is one of the main barriers to a cure of HIV/AIDS. Here, we show that holo-RBP4, the carrier of Vitamin A in the blood stream, reactivates latent HIV-1 in a variety of T cells lines as well as in a significant number of patientś derived cells. Holo-RBP4 is abundant in the circulation and has been reported to be further increased in PLWH under effective ART.^40^ We found that a surprisingly high number of hemofiltrate-derived fractions reactivates latent HIV-1 and this activity correlated with the presence of intact and C-terminally truncated RBP4 proteins. Notably, the relative concentrations of the RBP4 in the peptide/protein library correspond to those circulating in human blood^41^. In agreement with a relevant role of RBP4 as modulator of HIV-1 latency *in vivo,* significant reactivation from latency was observed at physiological concentrations of 10 to 50 µg/ml and confirmed using PBMCs obtained from PLWH after long-term cART and with undetectable viral RNA loads.

Accumulating evidence suggests that retinoic acid (RA), the biologically active metabolite of retinol, reactivates latent lentiviruses *in vitro* and *in vivo.*^42,43^ Initially, it has been shown that RA reactivates SIV in latently infected cells.^26^ More recently, it has been reported that RA also enhances activation of replication-competent HIV-1 reservoirs from subjects on suppressive cART.^42^ In addition, treatment with the RA derivative acitretin was shown to enhance RIG-I signaling and HIV-1 transcription.^44^. Thus, it came as surprise that we only observed minimal effects of RA treatment on HIV reactivation (Fig. 3d). This was not just due to inefficient uptake since treatment of J-Lat cells with agents preventing the conversion of retinal to retinoic acid upon delivery by holo-RBP4 did not inhibit RBP4-induced reactivation of latent HIV-1 proviruses (Fig. 3e). Altogether, our results indicate that RBP4-mediated delivery of retinol and subsequent oxidation to retinaldehyde (retinal) and retinoic acid (RA) is not required for reactivation of HIV-1 in latently infected cells.

It is known that RBP4 binding to its STRA6 receptor not only mediates delivery of retinol into the cells but also triggers the JAK/STAT5 (Janus kinase/signal transducer and activator of transcription) pathway.^31^ Notably, retinol is required for efficient binding of RBP4 to STRA6 and receptor-mediated signaling. This helps to explain why only holo-RBP4 efficiently reactivated latent HIV. In addition to JAK/STAT signaling, RBP4 has been reported to activate the c-Jun N-terminal protein kinase (JNK) pathway upon interaction with Toll-like receptor 4 (TLR4) independently of STRA6.^45^ Stimulation of JNK activates transcription factors like AP-1 (Activator Protein-1), which has been reported to synergize with NF-kB to activate latent HIV-1.^46^ We found that STAT5 and JNK inhibitors both inhibit RBP4-mediated reactivation of latent HIV-1 in J-Lat cells in a dose-dependent manner by up to 50% and 60%, respectively (Fig. 4). While more research is needed to fully define the underlying mechanisms, STRA6- and/or TLR2/4-mediated activation of JAK/STAT5 and JNK signaling clearly play relevant roles in RBP4-dependent activation of latent HIV.

One interesting question is whether circulating RBP4 affects the latent HIV reservoirs *in vivo*. While RBP4 is mainly produced in the liver, it is also an adipocytokine linked to obesity and insulin resistance.^47^ Obese children and adult individuals with obesity and type 2 diabetes have significantly higher RBP4 levels compared with lean healthy controls.^36^ Notably, the mechanisms involved in reactivation of latent HIV-1, i.e. TLR2/4-mediated activation of JNK signaling and STRA6-dependent activation of JAK/STAT, have also been proposed to contribute to the detrimental effects of RBP4 on insulin sensitivity.^29,31,45^ It is thus tempting to speculate that the latent HIV-1 reservoirs may be altered in obese PLWH. Notably, adipose tissue also serves as reservoir for HIV and inflammation may both enhance viral replication due to the activation of NF-kB and other transcription factors but also suppress viral spread due to the induction of antiviral factors. Although the plasma levels of RBP4 in PLWH were well sufficient to reactivate HIV-1 in latency models, they did not correlate with the proviral loads in blood (Figure 5A). Altogether, the effects of holo-RBP4 on HIV replication, latency and pathogenicity are most likely complex and clearly warrant further investigation.

In conclusion, our study shows that holo-RBP4 plays a previously unrecognized role in the activation of latent HIV-1 proviruses by mechanisms involving JAK/STAT5 and TLR4/JNK signaling but independently of RA. As the primary carrier protein for retinol in the bloodstream, RBP4 delivers retinol to a wide range of tissues, such as the thymus, spleen, and lymph nodes. Thus, holo-RBP4 may exert effects in lymphoid tissues, the major site of the latent reservoirs of HIV. Our findings indicate that holo-RBP4 provides a new target to attack the latent reservoirs of HIV-1. Our results also demonstrate that the screening of peptide/proteins libraries from body fluids allows to discover effective natural LRAs. Currently, we are using this approach to identify endogenous agents reactivating HIV-1 proviruses even in latency models that are poorly responsive to PMA, TNF-α and other known LRAs.

## MATERIALS AND METHODS

### Purification of active RBP4 from human hemofiltrate

Human hemofiltrate is a waste product of hemodialysis of patients with chronic renal failure. From 10.000 of liters of hemofiltrate (batch 050905) a peptide library was generated.^41^ Peptides were separated into 384 fractions based on charge (cation-exchange separation) and hydrophobicity (reversed-phase liquid chromatography). This hemofiltrate peptide library is a salt-free source of highly concentrated peptides and small proteins (< 30 kDa) which can be used to screen for antimicrobial and antiviral peptides^12^. A reporter cell assay was carried out with the addition of different hemofiltrate fractions. One additional round of chromatographic purification in combination with the bioassay was performed to obtain a pure active compound. For reversed-phase separation the Pool 4 fraction from the cation-exchange separation was subjected to reversed-phase chromatography on Source RPC 15 (Cytiva, USA) of dimensions 2 × 25 cm at a flow rate of 13 ml/min, using a linear gradient of acetonitrile using aqueous buffer A (0.1% TFA) and B (80% acetonitrile, 0.05%TFA) resulting in bioactive fraction P4, frs. 27-38. Detection of eluting compounds was online monitored at 280 nm. Using these bioactive fractions, the final purification step was performed on a Phenomenex Aqua RP18 (Phenomenex, USA) of dimensions 1 × 25 cm at a flow rate of 2.5 mL/min, using a linear gradient of acetonitrile using same RP buffers resulting in main bioactive fractions 34 and 42. Fraction 42, of higher purity was subjected to mass spectrometry and sequence analyses.

### Molecular mass measurement of RBP4 by MALDI-TOF MS

The sample was analysed using an Axima Confidence MALDI-TOF MS (Shimadzu, Japan) in positive linear mode on a 384-spot stainless-steel sample plate. Spots were coated with 1 µL 5 mg/mL CHCA previously dissolved in matrix diluent (Shimadzu, Japan), and the solvent was allowed to air dry. Then 0.5 µL sample or standard was applied onto the dry pre-coated well and immediately mixed with 0.5 µL matrix; the solvent was allowed to air dry. All spectra were acquired in the positive ion linear mode using a 337-nm N2 laser. Ions were accelerated from the source at 20 kV. A hundred profiles were acquired per sample, and 20 shots were accumulated per profile. The equipment was calibrated with a standard mixture provided in the TOFMixTM MALDI kit (Shimadzu, Japan). Measurements and MS data processing were controlled by the MALDI-MS Application Shimadzu Biotech Launchpad 2.9.8.1 (Shimadzu, Japan).

### Identification of RBP4 by LC-MS/MS sequencing

The sample was reduced with 5 mM DTT for 20 min at RT, then carbamidomethylated with 50 mM iodoacetamide for 20 min at 37 °C, and subsequently digested with trypsin (ThermoFisher Scientific, 900,589), at a 1:50 ratio (enzyme: protein) for 16 h at 37 °C. A 15 µl aliquot of the digested sample was analyzed using an Orbitrap Elite Hybrid mass spectrometry system (Thermo Fisher Scientific) online coupled to an U3000 RSLCnano (Thermo Fisher Scientific) uPLC as described ^48^. XCalibur 2.2 SP1.48 (Thermo Fisher Scientific, Bremen, Germany) was used for data-dependent tandem mass spectrometry (MS/MS) analyses. Database search (PEAKS DB) was performed using PEAKs X + studio. For peptide identification, MS/MS spectra were correlated with the UniProt human reference proteome set (Uniprot release 2023_03; 20,423 reviewed entries). Parent mass error tolerance and fragment mass error tolerance were set at 15 ppm and 0.5 Da, respectively. The maximal number of missed cleavages was set at 3. Carbamidomethylated cysteine was considered as a fixed modification, and methionine oxidation as a variable modification. False discovery rates were set on the peptide level to 1%.

### Cell culture

Latently HIV-1 infected cell lines were obtained from the NIH AIDS repository (*J-Lat full length cells*: J-Lat 6.3 #9846, J-Lat 8.4 #9847, J-Lat 9.2 #9848, J-Lat 10.6 #9849, J-Lat 15.4 #9850; *J-Lat GFP cells*: 82 #9851, A1 #9852, A7 #9853, A2 #9854, H2 #9855, A72 #9856;

*cells without integrated reporter gene*: Jurkat E6 J1.1 # 1340, U937 U1 #165, ACH-2 #ARP-349). Cells were cultivated in Roswell Park Memorial Institute (RPMI, Gibco) 1640 medium supplemented with 10 % (v/v) FBS (Gibco), 100 U/ml penicillin (PAN-Biotech), 100 µg/ml streptomycin (PAN-Biotech) and 2 mM L-glutamine (PAN-Biotech). Cells were tested for mycoplasma contamination by polymerase chain reaction (PCR) test and used if negative. Cells were split 1:10 regularly 2-3 times a week.

### Generation and screening of HF-libraries

Fractions of a hemofiltrate-derived peptide library generated as described before^12^ were tested for their ability to reactivate latent HIV-1 proviruses in J-Lat 11.1 cells. For this, 100,000 J-Lat cells were seeded in F-bottomed 96-well dishes and incubated with the peptide fractions for 24 h. The next day, cells were analysed for GFP expression by flow cytometry (s. section Flow cytometry for details)

### Activity of purified RBP4 on several latently infected cell lines

To investigate the effect of purified RBP4 on different latently infected cell lines, 300,000 cells of each cell line were seeded in 48-well dishes and treated with the indicated concentrations of plasma-derived, purified RBP4 (Medix Biochemica, #527-12-1) in duplicates. As positive controls, cells were treated with 5 ng/mL PMA or TNFα. Cells were incubated at 37°C, 90% humidity and 5% CO2. 2 days post treatment, cells were washed, fixed with 2% (v/v) PFA and analysed by flow cytometry (for detail see *Flow Cytometry*).

### Cell viability staining

Cell were spinned down at 350 x g and room temperature for 3 minutes. Supernatants were discarded. Cells were washed twice in 1x PBS. eBioscience™ Fixable Viability Dye eFluor™ 780 (Thermo Scientific, #65-0865-14) was diluted 1:1000 (v/v) in 1x PBS. Cells pellets were resuspended in 50 µL solution containing the viability dye and incubated at room temperature for 15 minutes in the dark. Cell were washed twice in 1x PBS. Finally, cells were fixed in 4% PFA at 4°C for 1 hour and analysed by flow cytometry.

### Flow Cytometry

All cells analysed by flow cytometry were gated based on forward and side scatter characteristics, followed by exclusion of doublets, followed by (optional: the viability dye positive and negative cells and) the analysis of GFP (J-Lat full length and J-lat GFP cells) or p24 (J1.1, U937, ACH-2 cells) expression. Data were generated with BD FACS Diva 6.1.3 Software using the FACS Canto II flow cytometer. Data analysis was performed using FlowJo 10.6 Software (Treestar) or FlowLogic Solution 1.0.

### Whole-cell lysates

Whole-cell lysates were prepared by collecting cells in Phosphate-Buffered Saline (PBS, Gibco). The cells were pelleted (500 x g, 4 °C, 5 min) and lysed in transmembrane lysis buffer (150 mM NaCl, 50 mM 4-(2-hydroxyethyl)-1-piperazineethanesulfonic acid (Hepes) pH 7.4, 1 % Triton X-100, 5 mM ethylenediaminetetraacetic acid (EDTA) by vortexing at maximum speed for 30 s. Cell debris were pelleted by centrifugation (20,000 x g, 4 °C, 20 min) and the cleared supernatants (SN) stored for analysis at -20 °C.

### SDS PAGE

Whole-cell lysates were mixed with 6xSDS-PAGE loading buffer (187.5 mM Tris-HCl pH 6.8, 75 % (v/v) Glycerol, 6 % (w/v) SDS, 0.3 % (w/v) Orange G, 15 % (v/v) 2-mercaptoethanol, modified from LI-COR) and heated to 95 °C for 5 min. Samples were loaded on a precast 10 % NuPAGE Bis-Tris gel (Invitrogen) and run for 2 h at 80 V in 1x MES SDS running buffer (Invitrogen). Subsequently, the gel was fixed with 50% MeOH/ 7% acetic acid for 15 min and washed 3 × 5 min with ultrapure water. Staining was performed with GelCode Blue Reagent (Thermo Fisher 24,590) overnight. The gel was destained using ultrapure water until the background appeared clear and then imaged in a LiCor Odyssey system.

### Treatment with inhibitors

300,000 J-Lat cells were seeded in 48-well dishes and treated with serial dilutions of 200, 400, 800 µM of DEAB (4-Diethylaminobenzaldehyde, Cat# HY-W016645, MedChemExpress) or 4, 8, 16 µM of WIN 18446 (Bis-diamine, Cat# SML3187, Sigma Aldrich) or 5, 10, 20, 40 µM of JNK inhibitor (JNK Inhibitor II, CAS 129-56-6 , Calbiochem) or 25, 50, 100 µM of STAT5 inhibitor (STAT5 Inhibitor, Cat# Cay15784-5, Biomol) or 50 µM of Rottlerin (Rottlerin, Cat# R5648, Sigma Aldrich) or 50 nM Selumetinib (AZD6244, Cat# HY-50706, MedChemExpress) or 50 µM of U0126 ( U0126, CAS 109511-58-2, Calbiochem) and incubated for 4 hours at 37°C. Cells were then treated with 5 ng/mL of TNFα (Tumor Necrosis Factor-α human, Cat# H8916, Sigma-Aldrich), 25 µg/mL, 50 µg/mL of RBP4 (Retinol Binding Protein 4, Cat#527-12-1, Medix Biochemica) and incubated at 37°C. 48 hours later, J-Lat cells were washed once with PBS and incubated with 100µL of eBioscience™ Fixable Viability Dye eFluor™ 780 diluted 1:1000 (v/v) in PBS for 15 minutes at RT in the dark. Cell were washed twice in PBS, fixed in 4% PFA at 4°C for 1 hour and analysed by flow cytometry.

### Treatment with retinoids

300,000 J-Lat cells were seeded in 48-well dishes and treated with serial dilutions of 1.5, 3, 6, 12.5 µg/mL of atRA (All-trans-Retinal, Cat# R2500, Sigma Aldrich) or 1.5, 3, 6, 12.5 µg/mL of RO (Retinol, Cat# R7632, Sigma Aldrich) or 0.75, 1.5, 3, 6, 12.5 µg/mL of RA (Retinoic Acid, Cat# R2625, Sigma Aldrich) or 25, 50, 100 µM of STAT5 inhibitor (STAT5 Inhibitor, Cat# Cay15784-5, Biomol) or 5 ng/mL of TNFα (Tumor Necrosis Factor-α human, Cat# H8916, Sigma-Aldrich), 100 µg/mL of Holo-RBP4 (Retinol Binding Protein 4, Cat#527-12-1, Medix Biochemica), 100 µg/mL of Apo-RBP4 (Recombinant Human RBP4 (carrier-free) Cat# 594906, Biolegend) and incubated at 37°C. 48 hours later, J-Lat cells were washed once with PBS and incubated with 100µL of eBioscience™ Fixable Viability Dye eFluor™ 780 diluted 1:1000 (v/v) in PBS for 15 minutes at RT in the dark. Cell were washed twice in PBS, fixed in 4% PFA at 4°C for 1 hour and analysed by flow cytometry.

### Primary CD4+ T cells

CD4 + T cells were isolated from buffy coats using the RosetteSep™ Human T Cell Enrichment Cocktail (Stem Cell, #15021) according to the manufacturer’s recommendation. CD4+ T cells were activated using Dynabeads Human T-Activator CD3/CD28 beads (Thermo Fisher), and cultured in RPMI-1640 medium containing 10% FCS, glutamine (2 mM), streptomycin (100 µg/ml), penicillin (100 U/ml) and interleukin 2 (IL-2, Miltenyi Biotec #130-097-745) (10 ng/ml).

### qRT-PCR

STRA6, IL-6 and IL-1β mRNA levels were determined in cells and normalized to GAPDH expression levels. Total RNA was isolated using the RNeasy Plus Mini kit (Qiagen) according to the manufacturer’s instructions. qRT-PCR was performed according to the manufacturer’s instructions using TaqMan Fast Virus 1-Step Master Mix (Thermo Fisher) and an OneStepPlus Real-Time PCR System (96-well format, fast mode). All reactions were run in duplicates using TaqMan primers/probes from Thermo Fisher: GAPDH (Cat# 4310884E), STRA6 (Hs00980261_g1), IL-1β (Hs01555410_m1), IL-6 (Hs00174131_m1).

### RBP4 ELISA

Concentration of RBP4 in plasma was measured using Human RBP4 ELISA Kit (Cat# DRB400, R&D) according to manufacturer’s instructions.

### HIV-1 Gag-p24 Single-Molecule Assay (Simoa)

Cryopreserved PBMCs from seven (7) HIV-infected ART-suppressed study participants were used to generate *ex vivo* measures of HIV-1 latency reversal as quantified by HIV-1 gag-p24 Single-Molecule Assay (Simoa) previously described^39,49^. Briefly, 20 million cells were cultured in 2 mL R10+ media plus 100 U/mL IL-2, 200 nM Raltegravir, in the presence of the test agent (RBP4) or anti-CD3/28 positive control and incubated for 72 hours at 37 °C and 5% CO_2_. By way of trypan blue stain, live cells were counted after incubation. Cell pellets and culture supernatants were harvested for viral gag-p24 protein Simoa. To harvest cells pellets for Simoa, we resuspended pellets in Simoa buffer (1x protease inhibitor cocktail, 49% FBS, 1% triton, 49% Blocker Casein in PBS (Thermo-Fisher)) and incubated for 30 minutes and frozen at -80°C. To harvest the culture supernatants, we added 10% triton (in PBS) to the supernatants to a final concentration of 1% and stored them at -80°. The samples were sent to University of Pennsylvania Human Immunology Core Facility and analyzed on the Quanterix HD-X machine using HIV p24 Quanterix kit (catalog 102215, lot 504201) according to the manufacturer protocol. Briefly, 278 µl of each sample and reagents were loaded to the plate and machine. The assay report was exported from the HD-X and the results were reported as pg/mL

### Intact Proviral DNA Assay (IPDA)

IPDA was used to assess the composition and levels of intact and defective proviruses in CD4+ T cells isolated from leukapheresis-derived PBMC as described^50^. Accelevir Diagnostics performed sample processing and IPDA quantification in accordance with the company standard guidelines by blinded operators. Briefly, isolation of the CD4+ T cells from cryopreserved PBMCs was conducted using EasySep Human CD4+ T cell Enrichment Kit (Stemcell Technologies). QIAamp DNA Mini Kit (Qiagen) was used to extract genomic DNA, whose concentration was then measured using fluorometry (Qubit dsDNA BR Assay Kit, Thermo Fisher Scientific). The quality was assessed using ultraviolet-visible (UV/VIS) spectrophotometry (QIAxpert, Qiagen). Finally, the genomic DNA was assessed by IPDA using the Bio-Rad QX200 AutoDG Droplet Digital PCR system. Results were expressed as count of total proviruses, intact, 3’ or 5’ deleted and/or hypermutated HIV pro-viruses per 106 CD4+ T cells.

### Statistical analysis

Results were graphed by Graph Pad Prism 10, and are displayed as means ± standard error of the mean (SEM). *P*-values ≤ 0.05 were considered statistically significant. A p-value below 0.05 is annotated with *, p-values below 0.01 with **, and p-values below 0.001 with ***.

### List of Supplementary Materials

Supplementary Figs. S1 to S6

## Supporting information

Six supplemental figures

## Acknowledgments

We thank Kenneth Lynn and Matthew Fair for their valuable assistance with patient-based studies. We also thank Shariq Usmani and Wolf-Georg Forssmann for their support and reagents during the early stage of this project. This study was supported by the DFG (CRC 1279), the NIH (AI164570, P30 AI045008), the Robert I. Jacobs Fund of the Philadelphia Foundation (LM), and a Herbert Kean, M.D., Family Professorship (LM). Additionally, we thank the Human Immunology Core (HIC) at the Perelman School of Medicine at the University of Pennsylvania for their support with the Simoa assay, with partial funding from NIH P30 AI045008 and P30 CA016520. The HIC is also supported by NIH grants and is identified by RRID: SCR_022380. G.M.L. was supported by NIH U24AI143502

## Author contributions

Conceptualization: KMJS, LM, FK; Methodology: CP, KR, MH, LK, AB, SU, NP, LS, AR, SW, EP, MF, GML and JM; Investigation: CP, KR, MH, LK, AB, SU, NP, LS, AR, SW, EP, MF and JM; Funding acquisition: KMJS, LM, FK; Supervision: KMJS, LM, FK; Writing – original draft: FK; Writing – review & editing: all authors

## Competing interests

Authors declare that they have no competing interests.

### Data and materials availability

All data are available in the main text or the supplementary materials. Materials are available from the corresponding author.

