## Supplementary material for "Retinol Binding Protein 4 reactivates latent HIV-1 via the JAK/STAT5 and JNK pathways": Six supplemental figures

[1] Institute of Molecular Virology, Ulm University Medical Centre, 89081 Ulm, Germany; [2] HIV Cure and Viral Diseases Center, The Wistar Institute, Philadelphia, PA 19104 USA; [3] Core Facility Functional Peptidomics (CFP), Ulm University Medical Center, 89081 Ulm, Germany; [4] Department of Pathology and Laboratory Medicine, Perelman School of Medicine, University of Pennsylvania, PA, 19104, USA; [5] Core Unit Mass Spectrometry and Proteomics (CUMP), Ulm University Medical Center, 89081 Ulm, Germany; [6] AccelevirDx, Baltimore, MD, USA, [7] German Center for Neurodegenerative Diseases (DZNE), Ulm, Germany

**This file includes:**

Supplementary figures S1 to S6

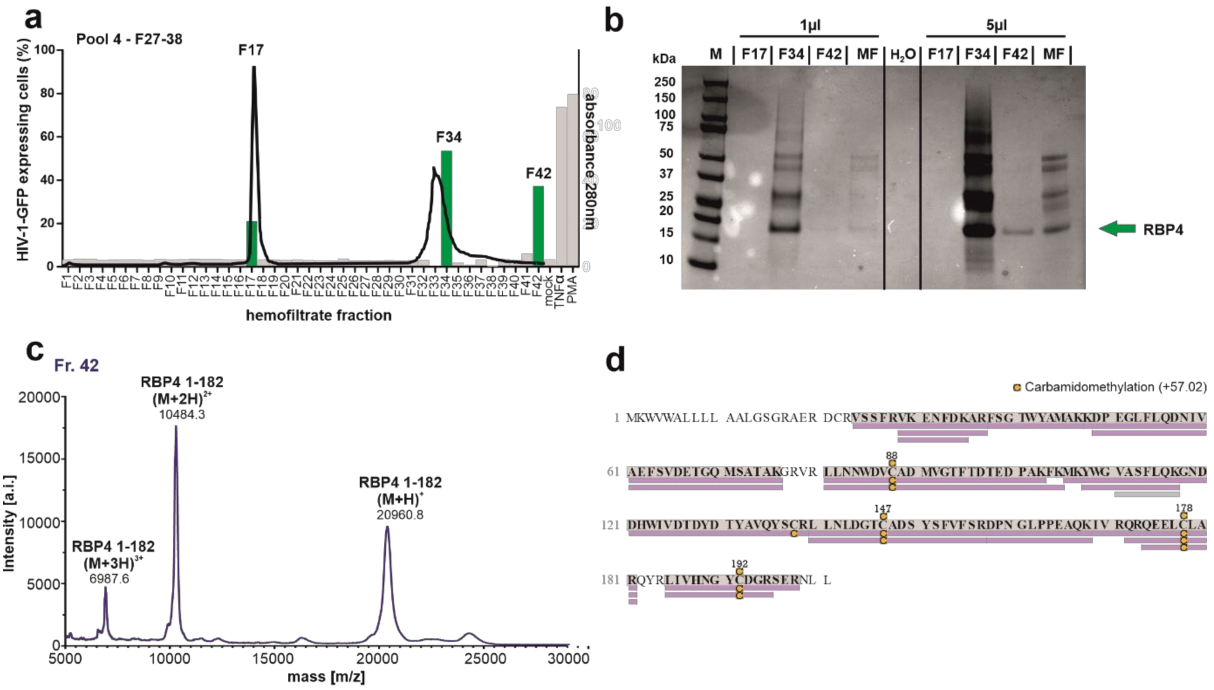


**Figure S1. Identification of RBP4 as activator of latent HIV-proviruses**. (**a**) Bars indicate the percentage of activated (eGFP positive) J-Lat 11.1 cells in presence of the peptide fractions from pool 4 of the hemofiltrate-derived library, showed in Figure 1a. The black line indicates the peptide/protein elution profile. Green bars indicate the fractions used for further purifications. Mock indicates absence of the peptide fractions. PMA and TNFα are used as positive controls. (**b**) Coomassie Brilliant Blue staining of a blot of the green HF-derived fractions from (a). The green arrow indicates the presence of a protein of around 21 kDa in most active fractions. (**c**) The MALDI-TOF spectrum of RBP4 shows a dominant signal of m/z 20960.75, which closely matches the theoretical m/z value of the RPB4 (1-182, 20959.42 Da) truncated variant lacking the C-terminal leucin. (**d**) Identification of RBP4. The protein was carbamidomethylated, digested with Trypsin, and analyzed with a nanoLC-Orbitrap Elite system. The analysis of the proteolytic fragments showed the presence of RBP4, with a sequence coverage of 91.7% with respect to the full-length mature polypeptide (RBP4 precursor 19-201).

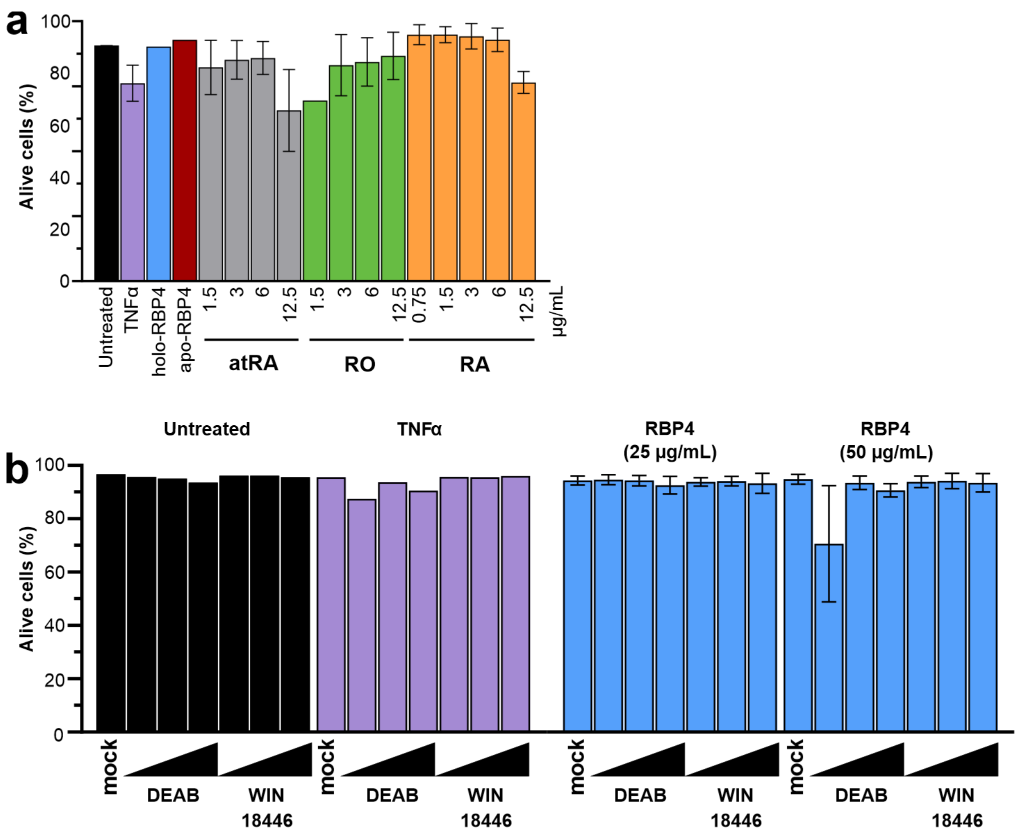


**Figure S2. Impact of RBP4 and free retinol and its metabolic products on cell viability.** (**a**) J-Lat 10.6 cells were treated with the indicated concentrations of all trans-retinal (atRA), retinol (RO) and retinoic acid (RA) or with TNFα, holo-RBP4 or apo-RBP4 and the percentage of alive cells were determined by flow cytometry. (**b**) J-Lat 10.6 cells were pre-treated with DEAB or WIN18,446 and subsequently left untreated or treated with TNFα or the indicated concentrations of RBP4. The percentage of alive cells was measured by flow cytometry.

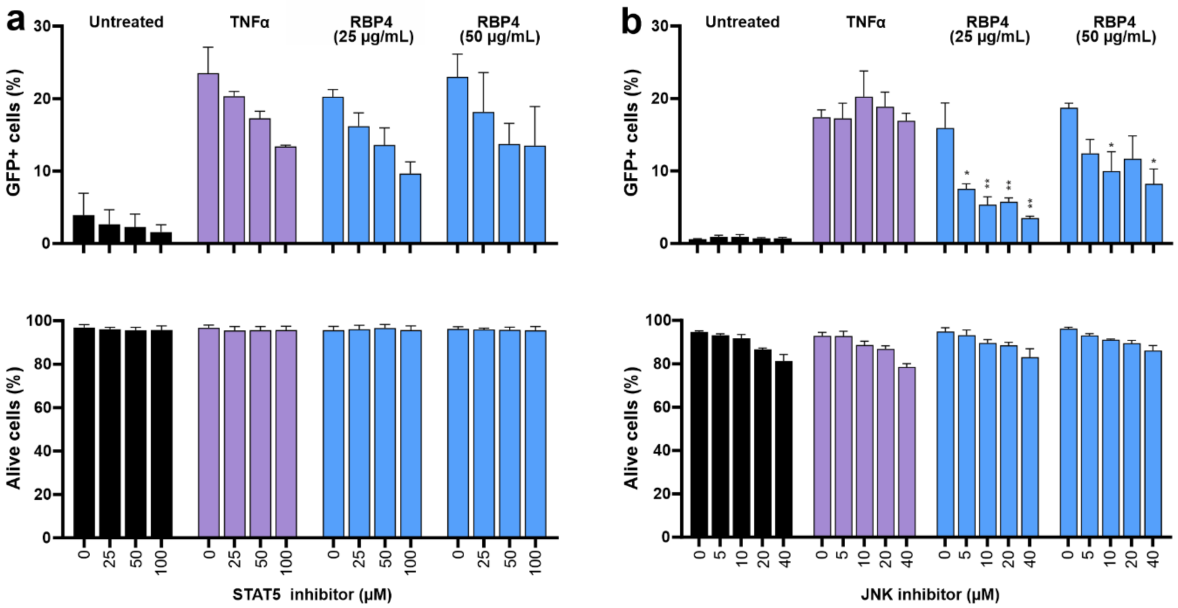


**Figure S3. Role of Jak/STAT and TLR/JNK activation in RBP4-mediated HIV reactivation.** (**a**)-(**b**) J-Lat 11.1 cells were pre-treated with the indicated concentrations of STAT5 (a) or JNK (b) inhibitor. 4 hours after, the cells were left untreated or treated with TNFα or the indicated concentrations of RBP4. The percentage of cells showing reactivation of HIV-1 eGFP reporter proviruses (upper panel) and the cell viability (lower panel) were determined by flow cytometry.

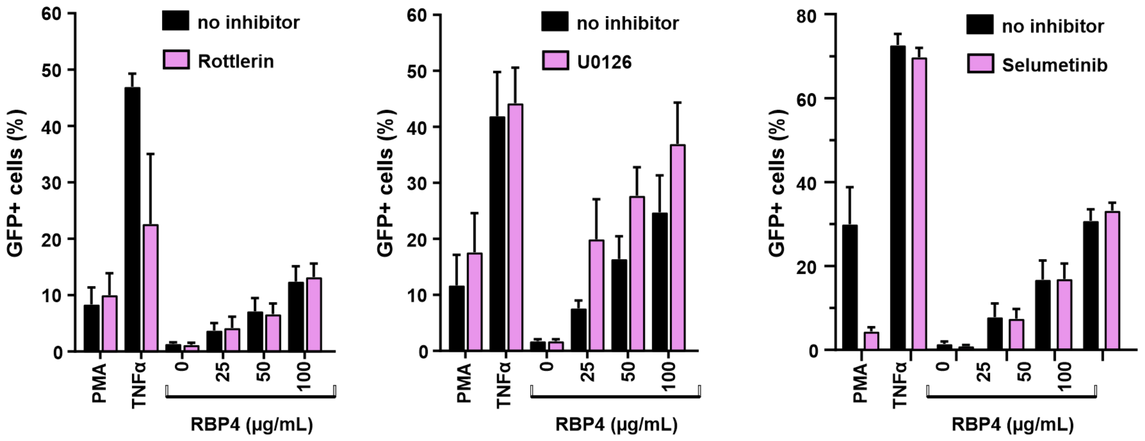


**Figure S4. Impact of PKC, MAPK and MEK1/2 on RBP4-mediated HIV reactivation**. (**a**)-(**b**)-(**c**) J-Lat 10.6 cells were left untreated (black bars) or pre-treated (pink bars) with the indicated concentrations of the PKC inhibitor Rottlerin (a) or the ERK1/2 inhibitor UO126 (b) or the MEK1/2 inhibitor Seleumetinib (c) .4 hour after, the cells were treated with PMA, TNFα or the indicated concentrations of RBP4. The percentage of cells showing reactivation of HIV-1 eGFP reporter proviruses were determined by flow cytometry.

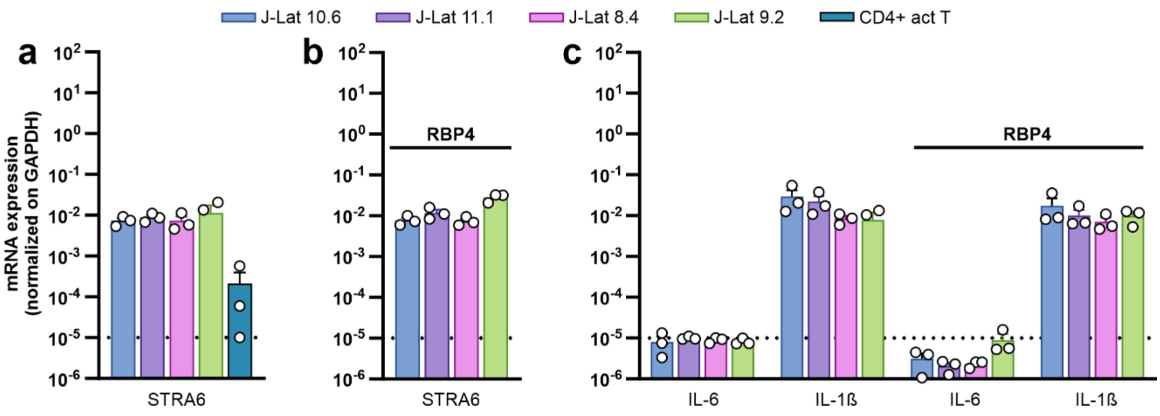


**Figure S5. STRA6, IL-6 and IL-1β expression in J-Lat cells.** Levels of STRA6 (**a**, **b**), IL-6 and IL-1β (**c**) mRNA expression in the indicated J-Lat cell lines or in primary CD4+ activated T cells that were left untreated or treated with 100 µg/mL of RBP4 (b). mRNA levels of the target genes were normalized on the GAPDH levels.

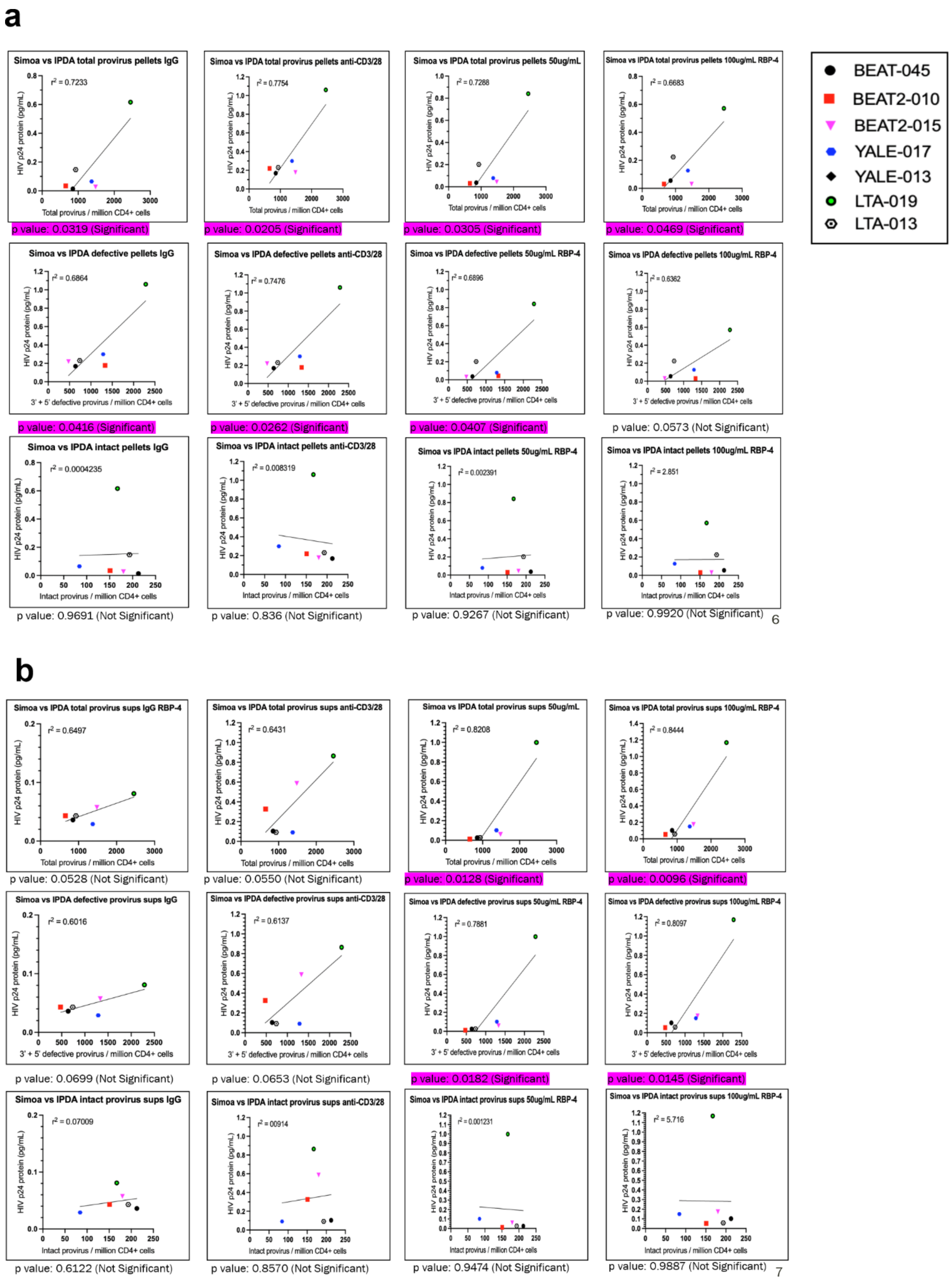


**Figure S6. Reactivation of latent HIV-1 in PBMCs from PLWH under cART.** (**a**)-(**b**) Correlation between total (upper panel) or defective (middle panel) or intact (lower panel) provirus/millions of CD4+ T cells from ART-suppressed PLWH and the HIV p24 induction in the cell pellets (a) or in the culture supernatants (b).
